# Rifampicin and rifabutin resistance in 1000 *Mycobacterium tuberculosis* clinical isolates

**DOI:** 10.1101/425652

**Authors:** Maha R Farhat, Jaimie Sixsmith, Roger Calderon, Nathan Hicks, Sarah Fortune, Megan Murray

**Affiliations:** Harvard Medical School, Department of Biomedical Informatics, Boston, MA; Massachusetts General Hospital, Division of Pulmonary and Critical Care, Boston, MA; Harvard Chan School of Public Health, Boston, MA; Socios en Salud, Lima, Peru; Harvard Medical School, Department of Global Health and Social Medicine, Boston, MA; Brigham and Women’s Hospital, Department of Global health and Equity, Boston, MA

## Abstract

Drug resistant tuberculosis (TB) remains a public health challenge with limited treatment options and high associated mortality. Rifamycins are among the most potent anti-TB drugs, and the loss of susceptibility to these agents, a hallmark of MDR TB, is considered a substantial therapeutic challenge. Rifamycins are known to target the RpoB subunit of RNA polymerase; however, our understanding of how rifamycin resistance is genetically encoded remains incomplete. Here we investigated *rpoB* genetic diversity and cross resistance between the two rifamycin drugs rifampicin (RIF) and rifabutin (RFB). We performed whole genome sequencing of 1005 MTB clinical isolates and measured minimum inhibitory concentration (MIC) to both agents on 7H10 agar using the indirect proportion method. Of the 1005 isolates, 767 were RIF resistant, and of these, 211 (27%) were sensitive to RFB at the critical concentration of 0.5ug/ml; 101/211 isolates had the *rpoB* mutation D435V (E.coli D516V). Isolates with discrepant resistance (RIF R and RFB S) 16.9 times more likely to harbor a D435V mutation as those resistant to both agents (OR 95% CI 10.5-27.9, P-value <10^-40^). To further understand this discrepancy, we generated both D435V and S450L (E.coli S531L) *rpoB* mutants in a laboratory strain and measured their antibiotic susceptibility using the alamar blue reduction assay. Compared with wildtype, D435V increased the 50% inhibitory concentration (IC50) to both RIF and RFB, however in both cases to a lesser degree than the S450L mutation. The observation that the *rpoB* D435V mutation produces an increase in the IC50 for both drugs contrasts with findings from previous smaller studies that suggested that isolates with D435V mutation remain RFB susceptible despite being RIF resistant. Our finding thus suggests that the recommended critical testing concentration for RFB should be revised.

## Introduction

Rifampicin resistance (RR) is the hallmark of multidrug resistance (MDR) in TB. By the WHO’s most recent estimates, 4.1% of new TB cases are RR/MDR and this rate is 19% among retreatment cases^1,2^. The estimated global incidence of RR/MDR was 600,000 in 2016, a figure that increased from 480,000 the year prior^1,2^. Timely and accurate diagnosis of RR remains a major challenge and together with challenges related to the duration and complexity of treatment regimens, progress in RR/MDR control has been limited. There is, however, increasing adoption of molecular diagnostic tests that rapidly detect genetic mutations in the RNA polymerase β subunit (*rpoB*) gene as a proxy for resistance to this drug class and MDR^2^. There are over 25 different resistance mutations detected by the latest generation of one such test the GeneXpert TB/RIF Ultra, and the results are currently summarized as binary ‘RIF resistance detected’ vs ‘non detected’^3^. At the same time, it is well recognized that genetic mutations can have a range of effects on the resistance phenotype, and that these effects can vary by bacterial lineage or genetic background^4,5^. This variability may extend to differential sensitivity to other drugs from the same class^6^. Given the complexity and side effects associated with MDR-TB therapy compared to susceptible TB, identifying subtypes of MTB isolates that are apparently resistant but may continue to be treatable with a higher dose of a first line agent or a related drug from the same class is highly desirable.

Whole genome sequencing (WGS) of MTB isolates is generating an abundance of information on the bacterial pathogen. To date most research utilizing WGS data has attempted to recapitulate the results of phenotypic culture-based drug susceptibility tests (DST) in the hope of WGS potentially replacing this expensive and time consuming approach^7–9^. However the power of WGS lies in potentially going beyond DST to allow the personalization of antibiotic therapy based on genotypic information. One such mutation, *rpoB* D435V thought to encode about 30% of RIF resistance, has been highlighted as associated with rifabutin (RFB) susceptibility at the CLSC recommended critical cutpoint of 0.5mg/L^10–12^. The drug RFB is considered to be of equal efficacy as RIF in drug susceptible TB, and it’s used in place of RIF in HIV patients with potentially interacting anti-retroviral therapy^13^. A few small or uncontrolled studies of RFB in patients with RR/MDR have suggested potential efficacy but there have been no prospective trials^14–16^. Most of the data has stemmed from comparisons between RIF & RFB DST and *rpoB* sequencing of MTB clinical isolates^17–22^. Constructing *rpoB* mutant alleles in isogenic laboratory strains can directly measure the effect of mutations like D435V on rifamycin cross resistance, but reports published to date have been conflicting. Williams et al. introduced *rpoB* D435V mutation through plasmid transduction into H37Rv lab strain and measured a step up in minimum inhibitory concentration (MIC) for RIF but not for RFB^23^. Gill and Garcia on the other hand measured a large step up in the rifamycin concentration needed to inhibit 50% of *ex-vivo* transcription with the introduction of the *rpoB* D435V for both RIF & RFB^24^. Notably the D435V mutation has been consistently found to have lower effects on the RIF MIC than S450L in several reports^4^. To try to add clarity to the debate, here we report on the largest collection of clinical MTB isolates where RIF & RFB resistance were quantified with MICs and compared with WGS results. We describe the diversity of rpoB variants seen, assess the degree of RIF/RFB cross resistance, and construct *rpoB* mutants in a laboratory strain to study the *in vitro* effect of D435V on RIF and RFB MIC.

## Patients and Methods

### Patient Cohort/Sample Collection

MTB sputum based culture isolates from Peru were selected to over represent MDR from samples collected for clinical care and then archived at the Massachusetts State Laboratory^7^ (n=496) or (2) sampled from a longitudinal cohort of patients with Tuberculosis from Lima Peru^25^ (n=568).

### Culture and Drug resistance/MIC testing

Lowenstein-Jenson (LJ) culture was performed from sputum specimens using standard NALC-NaOH decontamination. All isolates, underwent MIC testing at NJH for RIF and RFB on 7H10 agar media using the indirect proportion method in a staged fashion. Isolates were first tested at three low concentrations that include the WHO recommended critical concentration. If the isolate was resistant at the critical concentration then testing at six higher concentration was also performed. The testing concentrations deviated from the traditional doubling to better detect intermediate level MICs that are close to the clinical critical concentration and within theoretically achievable levels in patient sera based on available pharmacodynamics data^26^. The drug concentrations were as follows (in µg/mL): RIF low: 0.25, 0.5, 1 and high: 2, 3, 5, 8, 10, 50; for RFB low: 0.125, 0.25, 0.50 and high: 0.60, 0.75, 0.875, 1.0, 1.5, 2.5.

### DNA extraction and Whole genome sequencing

DNA from sputum samples of TB patients was extracted from cryopreserved cultures. Each isolate was thawed and subcultured on LJ and a big loop of colony growth was lysed with lysozyme and proteinase K to obtain DNA using CTAB / Chloroform extraction and ethanol precipitation. DNA was sheared into ~250bp fragments using a Covaris sonicator (Covaris,Inc.), and prepared using the TruSeq Whole-Genome Sequencing DNA sample preparation kit (Illumina, Inc.) and sequenced on an Illumina HiSeq 2500 sequencer with paired-end reads of length 125 bp.

### Variant calling and phylogeny construction

We aligned the Illumina reads to the reference MTB isolate H37Rv using Stampy 1.0.23^27^ and variants were called by Platypus 0.5.2^28^ using default parameters. Genome coverage was assessed using SAMtools 0.1.18^29^ and FastQC^30^ and read mapping taxonomy was assessed using Kraken^31^. Strains that failed sequencing at a coverage of less than 95% at ≥10x all sites in *rpoB* or had a mapping percentage of less than 90% to MTB complex were excluded. Variants were filtered if they had a quality of <15, purity of <0.4 or did not meet the PASS filter designation by Platypus. TB genetic lineage was called using the Coll *et al*.^32^ SNP barcode and confirmed by constructing a Neighbor joining phylogeny using MEGA-5 ^33^ including lineage representative MTB isolates from Sekizuka *et al*.^34^.

### Statistical analysis

The Fisher exact test was used for testing proportions and the Wilcoxon rank sum test was used for comparing MIC distribution using R version 3.2.3. The significance threshold was set at <0.01. A mutation was determined to be homoplasic if it was found in isolates that belonged to more than one sublineage as defined by the Coll *et al*.^32^ SNP barcode.

### Construction and susceptibility testing of rpoB mutants

Using the laboratory MTB strain HN878 we obtained *rpoB* mutants by isolating individual colonies from cultures selected on solid 7H10 (Middlebrook 7H10 media, 0.5% glycerol, 10% OADC, 0.05% Tween-80) media containing RIF at 2 mg/L as previously described^35^. The RIF resistance determining region (RRDR) of *rpoB* (codons 426 – 454) was sequenced for each colony and the causal mutation was identified. We measured the MIC of the parental HN878 strain, as well as D435V and S450L mutants against both RIF and RFB using the alamar blue reduction method. Briefly, strains were grown to mid logarithmic phase in 7H9 media (7H9 salts, 0.2% glycerol, 10% OADC, 0.05% Tween-80) and subcultured to OD_600_ 0.003 in 200ul of 7H9 media in 96-well plates containing either no antibiotics or varying concentrations of RIF and RFB. The concentrations tested in µg/mL were RIF: 32, 8, 2, 0.5, 0.125, 0.03, 0.008, 0.002 and RFB: 0.5, 0.125, 0.03, 0.008, 0.002, 0.0005, 0.00012, 0.00003. Each condition was performed in three independent triplicate wells. Strains were cultured for 4 days with shaking at 37C and then 1/10 volume of alamar blue (BioRad) was added. After 4 additional days of culture, the reduction of alamar blue was quantified by measurement of optical density at 570nm. Normalized growth was calculated for each replicate by subtracting the OD_570_ of cell free well from each condition, and scaling by the background-subtracted OD_570_ measurement of a no drug well. Because of relatively high background OD seen at low drug concentrations with the alamar blue reduction assay we report the fold increase in IC50 rather than the MIC.

## Results

Of the total number of isolates, 59 had sequencing data that failed quality criteria and were excluded. In the remaining 1005 isolates, 767 were RR (MIC>1mg/L), and of these 211 (27%) displayed a RFB MIC ≤0.5mg/L. Table 1 compares the RIF and RFB MICs for the isolates. We identified 94 different genetic variants in the *rpoB* gene or upstream intergenic region that occurred in one or more RR isolate (Supplementary Table 1). The gene body contained 74 unique non-synonymous single or double nucleotide substitutions and 8 indels at 51 different codons. The three most common *rpoB* RIF resistance determining region (RRDR) mutations occurred at codons S450L, D435V and H445Y in 440, 131 and 21 RR isolates respectively. Several non-RRDR *rpoB* mutations were homoplasic i.e. occurred in isolates belonging to more than one TB sub-lineage. There were 5 such non-synonymous mutations that occurred in 5 or more isolates. Four co-occured with one or more RRDR mutation, but the overlap was not complete for two: E250G and V170F. Only the mutation I491F was homoplasic and occurred in isolates with no RRDR mutation found. Five other nearby mutations: P454R, I480V, I488L, I491S and L494P occurred in 1-3 RR isolates each for a total of 8 isolates. Although these variants did not display any homoplasy, there were no co-occurring RRDR mutations in those isolates to explain RR (Supplementary Table 1).

**Table 1:** RIF vs RFB MICs for the 1005 isolates. All concentrations are in µg/ml.

| *RIF* | *RFB* |  |  |  |  |  |  |  |  |  |
| --- | --- | --- | --- | --- | --- | --- | --- | --- | --- | --- |
|  | ≤0.125 | 0.250 | 0.500 | 0.600 | 0.750 | 0.875 | 1.0 | 1.5 | 2.5 | >2.5 |
| ≤0.250 | 131 | 0 | 0 | 0 | 0 | 0 | 0 | 0 | 0 | 1 |
| 0.500 | 63 | 0 | 0 | 0 | 0 | 0 | 0 | 0 | 0 | 0 |
| 1.0 | 40 | 1 | 1 | 0 | 0 | 0 | 0 | 0 | 0 | 0 |
| 2.0 | 30 | 1 | 4 | 3 | 0 | 0 | 0 | 0 | 0 | 0 |
| 3.0 | 8 | 0 | 1 | 0 | 2 | 0 | 0 | 0 | 0 | 0 |
| 5.0 | 9 | 0 | 0 | 0 | 0 | 0 | 0 | 0 | 0 | 0 |
| 8.0 | 12 | 2 | 1 | 3 | 2 | 0 | 0 | 1 | 0 | 1 |
| 10.0 | 6 | 0 | 0 | 0 | 0 | 0 | 0 | 0 | 0 | 2 |
| 50.0 | 37 | 11 | 3 | 9 | 0 | 0 | 1 | 6 | 3 | 2 |
| >50 | 44 | 27 | 15 | 17 | 9 | 10 | 15 | 36 | 83 | 352 |

Of the 211 isolates with RFB sensitivity and RR, to be labeled ‘discordant isolates’, 102 isolates carried the *rpoB* mutation D435V. Four additional variants: S450L, H445Y, D435Y were observed in >5% (10) of the discordant isolates. Overall, the discordant isolates carried 31 different nonsynonymous *rpoB* variants (Table 2). We tested all the observed variants for association with the discrepant vs. the concordant (RIF and RFB resistant) phenotype, significantly associated mutations are shown in bold in Table 2. Discordant isolates were more likely to carry a D435V, D435Y, H445L or H445C mutation than those resistant to both agents. On the other hand, concordant isolates were more likely to carry an S450L or a H445Y mutation. We tested if the mutations associated with discordance resulted in lower RIF MICs as compared with S450L and this was the case for all four mutations (P-value <10^-15^ in each case) (Figure 1). We also tested if there is a difference in RFB MIC observed among isolates from the Beijing (2.2) vs the Euro-American sublineages (4, 4.1, 4.3) with D435V mutations and observed the median MIC among the Beijing isolates to be 0.375 (IQR 0.188-1.5 µg/ml) significantly higher than the RFB MIC for the Euro-American pool of isolates median MIC ≤0.125 (IQR ≤0.125-0.188µg/ml) P-value 6×10^-4^.

**Table 2:** *rpoB* variant alleles found in isolates with discordant rifamycin resistance (RFB MIC <0.5 µg/ml and RIF MIC>1µg/ml) vs. concordant rifamycin resistance. *A695T only occurred only in lineage 4.3 isolates and always with S450L (46 isolates) or D435V (1 isolate) and thus this association is likely confounded; A695T was not observed in RIF susceptible isolates. See Supplementary table 1 for more information on non-RRDR variants

| *H37Rv*<br>position | Nucleotide<br>change | Amino acid<br>change | Count in<br>discrepant<br>(n=211) | Count in<br>concordant<br>(n=556) | RIF MIC<br>(median) | RIF MIC (IQR) | RFB MIC<br>(median) | RFB MIC (IQR) | OR | 95% CI | P-value |
| --- | --- | --- | --- | --- | --- | --- | --- | --- | --- | --- | --- |
| **761110** | **A>T** | **D435V** | **102** | **29** | **>50** | **( 30 - 50 )** | **≤0.125** | **( ≤0.125 – 0.375 )** | **16.91** | **( 10.51 , 27.92 )** | **5E-41** |
| **761155** | **C>T** | **S450L** | **19** | **421** | **>50** | **( 50 - 50 )** | **>2.5** | **( 2 – >2.5 )** | **0.03** | **( 0.02 , 0.05 )** | **3E-67** |
| 761889 | G>C | V695L | 18 | 51 | >50 | ( 1.5 - 50 ) | 0.9375 | ( ≤0.125 – 2 ) | 0.92 | ( 0.49 , 1.66 ) | 0.9 |
| **761109** | **G>T** | **D435Y** | **12** | **3** | **3.25** | **( 1.5 - 50 )** | **≤0.125** | **( ≤0.125 – 0.328 )** | **11.08** | **( 2.95 , 61.82 )** | **3E-05** |
| **761140** | **A>T** | **H445L** | **12** | **0** | **6.5** | **(2.125 - 6.5)** | **≤0.125** | **( ≤0.125 – ≤0.125 )** | **Inf** | **( 7.65 , Inf )** | **1E-07** |
| **761139** | **CA>TG** | **H445C** | **7** | **0** | **30** | **( 17 - 30 )** | **≤0.125** | **( ≤0.125 – ≤0.125 )** | **Inf** | **( 3.88 , Inf )** | **1E-04** |
| 761161 | T>C | L452P | 5 | 4 | 4 | ( 1.5 - 40 ) | ≤0.125 | ( ≤0.125 – 0.9 ) | 3.34 | ( 0.71 , 17.02 ) | 0.07 |
| 760555 | A>G | E250G | 4 | 9 | 1.5 | ( ≤0.25 - 50 ) | ≤0.125 | ( ≤0.125 – >2.5 ) | 1.17 | ( 0.26 , 4.26 ) | 0.76 |
| 761277 | A>T | I491F | 4 | 2 | 3.25 | ( 1.75 – 5.875 ) | 0.25 | ( ≤0.125 – 0.506 ) | 5.34 | ( 0.76 , 59.46 ) | 0.05 |
| 761095 | T>G | L430R | 4 | 1 | >50 | ( 30 - 50 ) | 0.375 | ( 0.188 – 0.375 ) | 10.69 | ( 1.05 , 527.53 ) | 0.02 |
| 761128 | C>T | S441L | 3 | 0 | 30 | ( 30 - 30 ) | ≤0.125 | ( ≤0.125 – 0.156 ) | - | ( 1.09 , Inf ) | 0.02 |
| **761880** | **G>A** | **A692T\*** | **2** | **45** | **>50** | **( >50 - >50 )** | **2** | **( 1.25 – >2.5 )** | **0.11** | **( 0.01 , 0.42 )** | **6E-05** |
| 761154 | TC>CA | S450Q | 2 | 3 | >50 | ( >50 - >50 ) | 0.55 | ( 0.375 – 0.935 ) | 1.76 | ( 0.15 , 15.49 ) | 0.62 |
| 761101 | A>C | Q432P | 2 | 1 | >50 | ( >50 - >50 ) | 0.375 | ( 0.281 – 0.525 ) | 5.30 | ( 0.27 , 313.25 ) | 0.19 |
| 761149 | G>A | R448Q | 2 | 1 | >50 | ( >50 - >50 ) | 0.375 | ( 0.375 – 0.812 ) | 5.30 | ( 0.27 , 313.25 ) | 0.19 |
| 761109 | G>A | D435N | 2 | 0 | >50 | ( >50 - >50 ) | 0.375 | ( 0.375 – 0.375 ) | - | ( 0.50 , Inf ) | 0.08 |
| 761148 | CG>AA | R448K | 2 | 0 | >50 | ( >50 - >50 ) | 0.1875 | ( 0.188 – 0.188 ) | - | ( 0.50 , Inf ) | 0.08 |
| **761139** | **C>T** | **H445Y** | **1** | **20** | **>50** | **( >50 - >50 )** | **>2.5** | **( >2.5 – >2.5 )** | **0.13** | **( 0.00 , 0.81 )** | **0.01** |
| 761139 | C>G | H445D | 1 | 7 | >50 | ( 45 - >50 ) | >2.5 | ( >2.5 – >2.5 ) | 0.37 | ( 0.01 , 2.94 ) | 0.46 |
| 760425 | G>A | E207K | 1 | 6 | >50 | ( >50 - >50 ) | >2.5 | ( >2.5 – >2.5 ) | 0.44 | ( 0.01 , 3.63 ) | 0.68 |
| 760645 | C>T | P280L | 1 | 0 | 30 | - | ≤0.125 | - | - | - | 0.28 |
| 761083 | delGCACCA | 426 | 1 | 0 | 6.5 | - | ≤0.125 | - | - | - | 0.28 |
| 761103 | insTTC | 433 | 1 | 0 | 30 | - | 0.1875 | - | - | - | 0.28 |
| 761108 | GGA>AGG | MD434-435IG | 1 | 0 | >50 | - | ≤0.125 | - | - | - | 0.28 |
| 761109 | GA>TT | D435F | 1 | 0 | 6.5 | - | ≤0.125 | - | - | - | 0.28 |
| 761115 | delAAC | 437 | 1 | 0 | >50 | - | ≤0.125 | - | - | - | 0.28 |
| 761139 | CA>GG | H445G | 1 | 0 | >50 | - | ≤0.125 | - | - | - | 0.28 |
| 761139 | CA>TC | H445S | 1 | 0 | 1.5 | - | ≤0.125 | - | - | - | 0.28 |
| 761167 | C>T | P454L | 1 | 0 | ≤0.25 | - | ≤0.125 | - | - | - | 0.28 |
| 761248 | A>C | E481A | 1 | 0 | 30 | - | 0.375 | - | - | - | 0.28 |
| 761905 | C>T | P700L | 1 | 0 | >50 | - | ≤0.125 | - | - | - | 0.28 |

**Figure 1:**
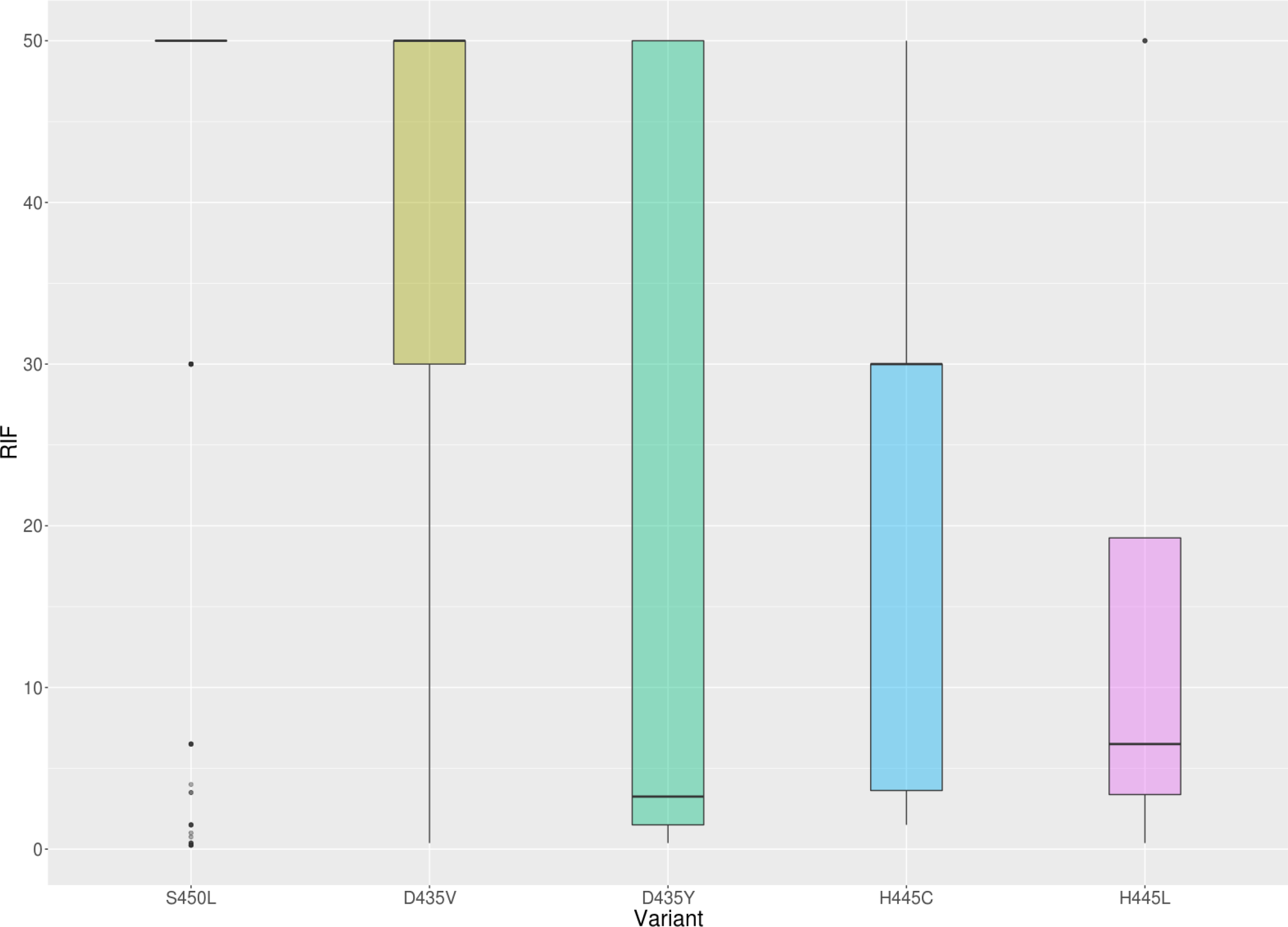
RIF MIC is significantly lower for the four mutations found to be associated with RIF-RFB discordance relative to S450L (associated with concordant resistance).

As D435V was the most common mutation associated with the apparent lack of cross resistance to RFB we selected a D435V and an S450L mutant in the laboratory strain HN878 by identifying spontaneous mutants resistant to RIF. Compared with the isogenic parental HN878 strain, the D435V mutation resulted in an increase in MIC against both RIF and RFB, however in both cases, this increase was less than what was observed for the S450L mutant (Figure 2). For RFB and D435V the IC50 was 62.5x that of wild type, as compared with >250x for S450L, for RIF D435V and S450L had IC50s that were 1000x and 16,000x of wild type respectively. The D435V mutant was almost completely inhibited at 0.5 µg/ml of RFB while S450L mutants retain substantial capacity for growth at that drug concentration.

**Figure 2:**
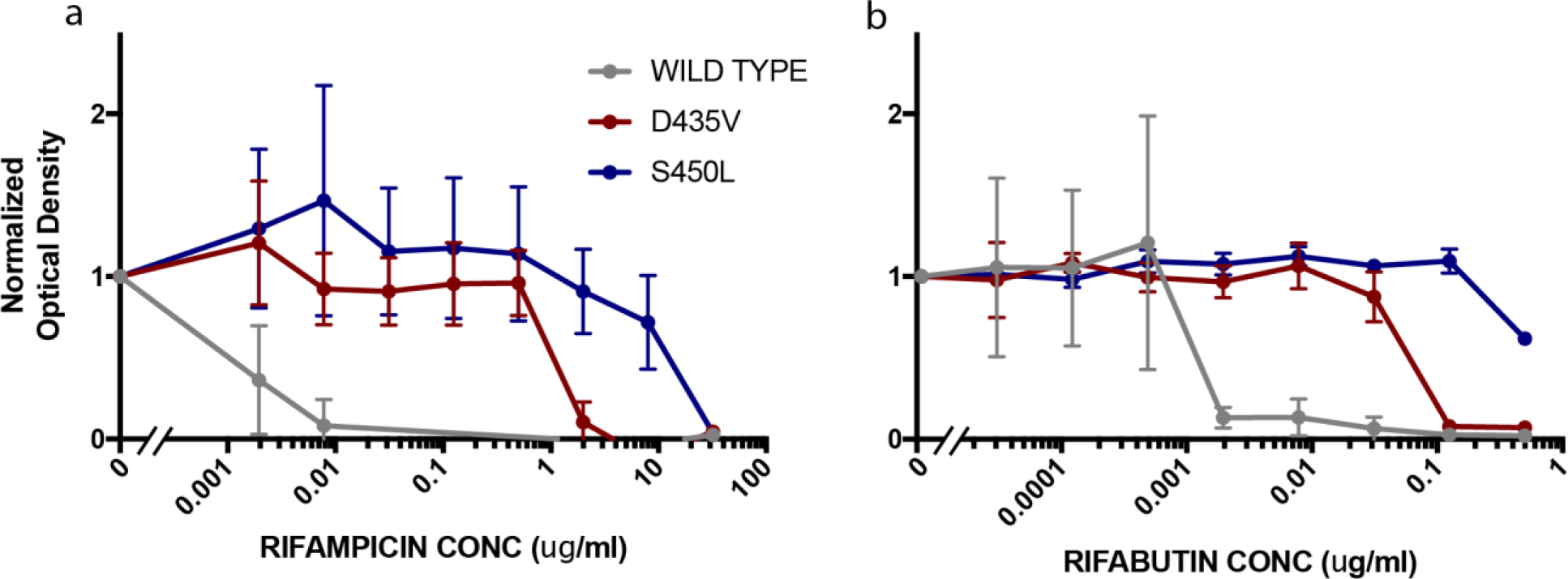
*rpoB* D435V mutations increase resistance to both RIF and RFB. Measurement of antibiotic resistance by alamar blue reduction assay in varying concentrations of RIF (a) and RFB (b). Optical density at 570nm is normalized with the growth of an antibiotic-free control set to 1. Each datapoint represents the mean of three independent cultures with the standard deviation.

## Discussion

In this study we observed RR isolates to have considerable *rpoB* genetic diversity. Several recent reports have challenged the notion that RR can only be caused by mutations within the RRDR, for example the I491F mutation was found to be a common cause of RR in Swaziland^36^. Homoplasy is one method of measuring variants under positive selection and hence of relevance to the resistance phenotype^37^. Although our study does not establish causation, we find several mutations outside of the RRDR, several in the vicinity of I491F, in RR isolates that have no other clearly causative mutation. This highlights the complexity of resistance even to the first drug rifampicin, and emphasizes the need to look beyond the RRDR, especially when patient treatment response is not as expected.

We also find that the lack of cross resistance between RIF and RFB to be relatively common, occurring in 27% of RR isolates. RFB resistance among RIF susceptible isolates was very rare and occurred in only one isolate. Again there was considerable *rpoB* variant diversity among the discordant isolates but the D435V mutation was by far the most common and occurred in nearly half of the discordant isolates (102/211, or 48%). Four mutations were significantly associated with discordance and two were significantly associated with concordance. Interestingly all four of the mutations significantly associated with discordance, were also associated with lower MICs to RIF itself and not just to RFB. This was also confirmed by measuring MICs in the two isogenic strains harboring D435V and S450L, with D435V found to raise both RIF and RFB MICs, albeit to a lower extent than S450L. The effect of D435V on RIF MIC does appear more profound than to RFB increasing the IC50 by 1000 fold vs 62.5 fold respectively, but we found D435V to increase IC50 more than has been previously reported in clinical isolates estimated at 2-4 fold of wild type MIC in one study that examined clinical isolates from the atypical Beijing lineage^10^. As HN878 belongs to the typical Beijing lineage we cannot exclude that the observed difference in inhibitory concentration may be a result of interaction between the D435V mutation and the genetic background. In the clinical isolate pooled examined, we did observe a higher RFB MIC in Beijing isolates than Euro-American isolates providing some evidence for interactions between the genetic background and resistance mutation.

Overall our finding that the D435V variant results in a large step up in RFB inhibitory concentration, especially in HN878 and East Asian-Beijing 2.2 isolates, suggests that RFB may have lower efficacy against such isolates, and that the RFB critical testing concentration, currently recommended at 0.5ug/ml may be too high. Rifamycin killing of MTB is concentration dependent, but achieving an adequate RFB peak serum concentration (Cmax ≥ 0.45µg/ml^38^) has proven challenging even when treating rifamycin sensitive TB^26,39^. Thus even small increases in RFB MIC may have significant consequences on treatment efficacy^26,39^. Ideally more retrospective data can be collected on the RFB treatment response of patients whose MTB isolates harbor mutations associated with discordance as there may not be equipoise to conduct a prospective RFB trial in such patients. Alternatively trials in which RFB is added to an otherwise adequate MDR region in patients with such isolates can be considered.

## Acknowledgments

We thank the Peruvian team for their patient care and for providing the patient isolates that enabled this research.

## Transparency

No relevant conflicts of interest to report for any of the coauthors.

## Data & Reproducibility

All sequence data is available on NCBI under Bioproject number PRJEB26000.

## References

1. World Health Organization. Global Tuberculosis Report 2016. (World Health Organization, 2016).

2. WHO | Global tuberculosis report 2017. Available at: http://www.who.int/tb/publications/global_report/en/. (Accessed: 1st August 2018)

3. Chakravorty, S. et al. The New Xpert MTB/RIF Ultra: Improving Detection of Mycobacterium tuberculosis and Resistance to Rifampin in an Assay Suitable for Point-of-Care Testing. mBio 8, e00812-17 (2017).

4. Nebenzahl-Guimaraes, H., Jacobson, K. R., Farhat, M. R. & Murray, M. B. Systematic review of allelic exchange experiments aimed at identifying mutations that confer drug resistance in Mycobacterium tuberculosis. J. Antimicrob. Chemother. (2013). doi:10.1093/jac/dkt358

5. Safi, H. et al. Evolution of high-level ethambutol-resistant tuberculosis through interacting mutations in decaprenylphosphoryl-β-D-arabinose biosynthetic and utilization pathway genes. Nat. Genet. (2013). doi:10.1038/ng.2743

6. Farhat, M. R. et al. Gyrase Mutations Are Associated with Variable Levels of Fluoroquinolone Resistance in Mycobacterium tuberculosis. J. Clin. Microbiol. 54, 727–733 (2016).

7. Farhat, M. R. et al. Genetic Determinants of Drug Resistance in Mycobacterium tuberculosis and Their Diagnostic Value. Am. J. Respir. Crit. Care Med. (2016). doi:10.1164/rccm.201510-2091OC

8. Bradley, P. et al. Rapid antibiotic-resistance predictions from genome sequence data for Staphylococcus aureus and Mycobacterium tuberculosis. Nature Communications 6, (2015).

9. Walker, T. M. et al. Whole-genome sequencing for prediction of Mycobacterium tuberculosis drug susceptibility and resistance: a retrospective cohort study. Lancet Infect Dis (2015). doi:10.1016/S1473-3099(15)00062-6

10. Sirgel, F. A. et al. The rationale for using rifabutin in the treatment of MDR and XDR tuberculosis outbreaks. PLoS ONE 8, e59414 (2013).

11. CLSI document M24-A. CLSI: Susceptibility Testing of Mycobacteria, Nocardiae, and Other Aerobic Actinomycetes; Approved Standard. (CLSI, 2003).

12. Dheda, K. et al. Outcomes, infectiousness, and transmission dynamics of patients with extensively drug-resistant tuberculosis and home-discharged patients with programmatically incurable tuberculosis: a prospective cohort study. The Lancet Respiratory Medicine 5, 269–281 (2017).

13. Official American Thoracic Society/Centers for Disease Control and Prevention/Infectious Diseases Society of America Clinical Practice Guidelines:… - PubMed - NCBI. Available at: https://www.ncbi.nlm.nih.gov/pubmed/27516382. (Accessed: 2nd August 2018)

14. Jo, K.-W. et al. The efficacy of rifabutin for rifabutin-susceptible, multidrug-resistant tuberculosis. Respir Med 107, 292–297 (2013).

15. Grassi, C. & Peona, V. Use of rifabutin in the treatment of pulmonary tuberculosis. Clin. Infect. Dis. 22 Suppl 1, S50-54 (1996).

16. Lee, H. et al. Treatment outcomes of rifabutin-containing regimens for rifabutin-sensitive multidrug-resistant pulmonary tuberculosis. Int. J. Infect. Dis. 65, 135–141 (2017).

17. Jamieson, F. B. et al. Profiling of rpoB Mutations and MICs for Rifampin and Rifabutin in Mycobacterium tuberculosis. J Clin Microbiol 52, 2157–2162 (2014).

18. Berrada, Z. L. et al. Rifabutin and Rifampin Resistance Levels and Associated rpoB Mutations in Clinical Isolates of Mycobacterium tuberculosis Complex. Diagn Microbiol Infect Dis 85, 177–181 (2016).

19. Chien, H. P., Yu, M. C., Ong, T. F., Lin, T. P. & Luh, K. T. In vitro activity of rifabutin and rifampin against clinical isolates of Mycobacterium tuberculosis in Taiwan. J. Formos. Med. Assoc. 99, 408–411 (2000).

20. Dickinson, J. M. & Mitchison, D. A. In vitro activity of new rifamycins against rifampicin-resistant M. tuberculosis and MAIS-complex mycobacteria. Tubercle 68, 177–182 (1987).

21. Senol, G., Erbaycu, A. & Ozsöz, A. Incidence of cross resistance between rifampicin and rifabutin in Mycobacterium tuberculosis strains in Izmir, Turkey. J Chemother 17, 380–384 (2005).

22. Uzun, M., Erturan, Z. & Anğ, O. Investigation of cross-resistance between rifampin and rifabutin in Mycobacterium tuberculosis complex strains. Int. J. Tuberc. Lung Dis. 6, 164–165 (2002).

23. Williams, D. L. et al. Contribution of rpoB Mutations to Development of Rifamycin Cross-Resistance in Mycobacterium tuberculosis. Antimicrob. Agents Chemother. 42, 1853–1857 (1998).

24. Gill, S. K. & Garcia, G. A. Rifamycin inhibition of WT and Rif-resistant Mycobacterium tuberculosis and Escherichia coli RNA polymerases in vitro. Tuberculosis (Edinb) 91, 361–369 (2011).

25. Zelner, J. et al. Protective effects of household-based TB interventions are robust to neighbourhood-level variation in exposure risk in Lima, Peru: a model-based analysis. International Journal of Epidemiology 47, 185–192 (2018).

26. Alsultan, A. & Peloquin, C. A. Therapeutic Drug Monitoring in the Treatment of Tuberculosis: An Update. Drugs 74, 839–854 (2014).

27. Lunter, G. & Goodson, M. Stampy: A statistical algorithm for sensitive and fast mapping of Illumina sequence reads. Genome Research 21, 936–939 (2011).

28. Rimmer, A. et al. Integrating mapping-, assembly- and haplotype-based approaches for calling variants in clinical sequencing applications. Nature Genetics 46, 912–918 (2014).

29. Li, H. et al. The Sequence Alignment/Map format and SAMtools. Bioinformatics (Oxford, England) 25, 2078–2079 (2009).

30. Babraham Bioinformatics - FastQC A Quality Control tool for High Throughput Sequence Data. Available at: https://www.bioinformatics.babraham.ac.uk/projects/fastqc/. (Accessed: 6th March 2018)

31. Wood, D. E. & Salzberg, S. L. Kraken: Ultrafast metagenomic sequence classification using exact alignments. Genome Biology 15, (2014).

32. Coll, F. et al. A robust SNP barcode for typing Mycobacterium tuberculosis complex strains. Nat Commun 5, 4812 (2014).

33. Tamura, K. et al. MEGA5: molecular evolutionary genetics analysis using maximum likelihood, evolutionary distance, and maximum parsimony methods. Mol. Biol. Evol. 28, 2731–2739 (2011).

34. Sekizuka, T. et al. TGS-TB: Total Genotyping Solution for Mycobacterium tuberculosis Using Short-Read Whole-Genome Sequencing. PLOS ONE 10, e0142951 (2015).

35. Ford, C. B. et al. Use of whole genome sequencing to estimate the mutation rate of Mycobacterium tuberculosis during latent infection. Nature Genetics 43, 482–486 (2011).

36. Sanchez-Padilla, E. et al. Detection of Drug-Resistant Tuberculosis by Xpert MTB/RIF in Swaziland. New England Journal of Medicine 372, 1181–1182 (2015).

37. Farhat, M. R. et al. Genomic analysis identifies targets of convergent positive selection in drug-resistant Mycobacterium tuberculosis. Nat. Genet. (2013). doi:10.1038/ng.2747

38. Weiner, M. et al. Association between acquired rifamycin resistance and the pharmacokinetics of rifabutin and isoniazid among patients with HIV and tuberculosis. Clin. Infect. Dis. 40, 1481–1491 (2005).

39. Boulanger, C. et al. Pharmacokinetic evaluation of rifabutin in combination with lopinavir-ritonavir in patients with HIV infection and active tuberculosis. Clin. Infect. Dis. 49, 1305–1311 (2009).

